# Nociceptor translational profiling reveals the RagA-mTORC1 network as a critical generator of neuropathic pain

**DOI:** 10.1101/336784

**Authors:** Salim Megat, Pradipta R. Ray, Jamie K. Moy, Tzu-Fang Lou, Paulino Barragan-Iglesias, Yan Li, Grishma Pradhan, Andi Wangzhou, Ayesha Ahmad, Robert Y. North, Patrick M. Dougherty, Arkady Khoutorsky, Nahum Sonenberg, Kevin R. Webster, Gregory Dussor, Zachary T. Campbell, Theodore J. Price

## Abstract

Pain sensing neurons, nociceptors, are key drivers of neuropathic pain. We used translating ribosome affinity purification (TRAP) to comprehensively characterize up-and down-regulated mRNA translation in *Scn10a*-positive nociceptors in chemotherapy-induced neuropathic pain. We provide evidence that an underlying mechanism driving these changes in gene expression is a sustained mTORC1 activation driven by MNK1-eIF4E signaling. RagA, a GTPase controlling mTORC1 activity, is identified as a novel target of MNK1-eIF4E signaling, demonstrating a new link between these distinct signaling pathways. Neuropathic pain and RagA translation are strongly attenuated by genetic ablation of eIF4E phosphorylation, MNK1 elimination or treatment with the MNK inhibitor eFT508. We reveal a novel translational circuit for the genesis of neuropathic pain with important implications for next generation neuropathic pain therapeutics.

**One Sentence Summary:** Cell-specific sequencing of translating mRNAs elucidates signaling pathology that can be targeted to reverse neuropathic pain

## Main text

Neuropathic pain is a devastating disease with extraordinarily poor treatment outcomes. Existing first-line treatments produce only 30% pain relief in the majority of patients (*1*). Chemotherapy-induced peripheral neuropathy (CIPN) is the primary dose-limiting side effect of cancer treatment and no drugs are approved to treat this form of neuropathic pain (*2*). Peripheral pain sensing neurons, called nociceptors, are the cellular origin of pain caused by neuropathies (*3*). The dynamics of nociceptors gene expression throughout the development of painful neuropathy at the genome scale is opaque. We employed the translating ribosome affinity purification (TRAP) technology (*4*), using the Nav1.8^Cre^ mouse to achieve sensory neuron-specific ribosome tagging with enrichment in the nociceptor population. We used these Nav1.8-TRAP mice to identify transcripts associated with ribosomes specifically expressed in nociceptors isolated from animals with or without CIPN. This tool elucidates pathology in a broad signaling network that centers around RagA and mechanistic target of rapamycin complex 1 (mTORC1) signaling that plays a central role in development of neuropathic pain. Our work provides new insights into the complex interplay of lysosomal mTORC1 signaling as a disease-causing factor in peripheral neuropathy. We demonstrate how this information provides new strategies for development of the first disease-modifying agents for neuropathic pain treatment, a debilitating and pervasive clinical problem.

To generate nociceptor-enriched ribosome-tagged mice, Nav1.8^cre^ mice were crossed with Rosa26^fs-TRAP^ (*5*), to express eGFP fused to the ribosomal L10a protein in Nav1.8-positive neurons. To ensure the specificity of our approach, we characterized expression of the transgene in the DRG **(Fig. 1A)**. We found that eGFP-L10a-positive neurons primarily co-localized with small diameter peripherin-positive neurons (81% ±3.1) while a smaller proportion of eGFP-labelled neurons co-expressed NF200, a marker of large diameter neurons (40.1% ± 2.8) **(Fig. 1B,C)**. The percentage of eGFP-L10a-positive neurons was equally distributed between non-peptidergic (86.7% ± 2.3) and peptidergic neurons (78.7%± 2.7) **(Fig.1C)**. We did not observe eGFP-L10a expression in non-neuronal cells in the DRG. Thus, this approach specifically labels sensory neurons including a substantial subset of neurons that are involved in nociception.

**Figure 1.**
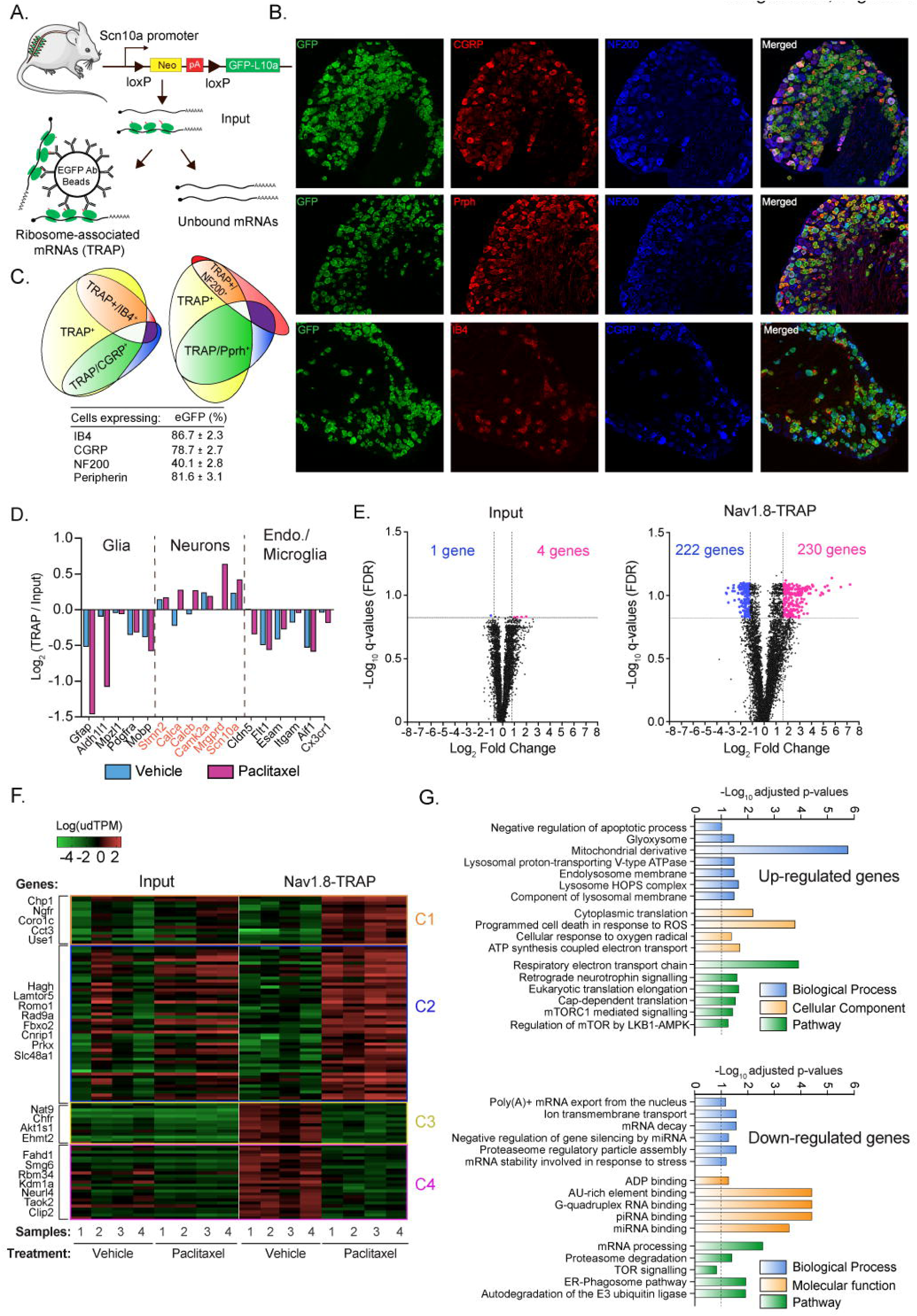
TRAP-seq identifies differentially translated mRNAs in Nav1.8-positive nociceptors. **(A)** Schematic representation of TRAP shows isolation of translating ribosomes (IP) from Input using anti-eGFP-coated beads. **(B-C)** Immunostaining for CGRP, IB4, NF200, and TRPV1 on L4-DRG sections from Nav1.8^cre^ / TRAP^floxed^ mice. A total of 9528, 8258 and 8526 neurons were counted for CGRP, NF200 and Prph staining, n = 3 mice/group. Scale bars, 100 μm. **(D)** DRG-TRAP-seq is enriched in neuronal markers such as *Calca* (CGRP) or *Camk2a* while glial (*Aldh1l1, Gfap*), endothelial (*Cldn5, Fit1*) and microglial (*Cxcr3, Aif1*) markers are depleted. **(E)** Volcano plot reveals 4 genes up-regulated and 1down-regulated in the transcriptome while more widespread changes are observed in the translatome (TRAP). **(F)** Heatmap shows the fold change of differentially expressed genes in each biological replicate of vehicle-and paclitaxel-treated animals (TRAP and Input fractions). **(G)** EnrichR analysis of the up-regulated genes in paclitaxel TRAP-seq samples.

The Nav1.8-TRAP was used to characterize the translatome of DRG neurons that express Nav1.8 using mRNAs isolated with immunoprecipitation (IP) of translating ribosomes (TRAP-seq) (*6*). First, we determined that IP of the eGFP-L10a from DRGs of control and Nav1.8-TRAP mouse strains yielded a specific IP of eGFP-L10a using eGFP antibodies **(Fig. S1A).** We also used IP combined with RT-qPCR for several mRNAs that are known to be enriched in nociceptors (*7*) to demonstrate integrity of the IP-mRNA complex **(Fig. S1B-C-D)**. These experiments demonstrate that the Nav1.8-TRAP approach can be used to tag ribosomes and capture mRNAs from nociceptive DRG neurons.

A large amount of starting material is required to successfully isolate ribosome-associated mRNAs from Nav1.8-TRAP cells. We determined that DRGs from cervical, thoracic and lumbar levels from 4 animals were minimally required for a single biological replicate of accurate TRAP-seq in this tissue. Since a goal of our experiments was to assess the nociceptor translatome in neuropathic pain, this prompted us to utilize a neuropathy model where all DRGs are affected such as CIPN. Almost 70% of cancer patients report symptoms of peripheral neuropathy during chemotherapeutic treatment and this is the primary dose-limiting side effect of chemotherapy treatment (*8*). Notably, the pathophysiology of CIPN, which is the most common cause of termination of chemotherapy (*2*) remains unknown.

We treated Nav1.8-TRAP mice with the commonly used chemotherapeutic agent, paclitaxel (4 mg/kg every other day, intraperitoneally, for a total of 8 days), which causes a robust small and large-fiber neuropathy in rodents and humans. To first determine whether paclitaxel increases the excitability of nociceptors at all vertebral levels we isolated cervical, thoracic and lumbar DRGs from mice treated with vehicle or paclitaxel and performed patch clamp electrophysiology. Paclitaxel treatment caused spontaneous activity and increased nociceptor excitability **(Fig. S2A-C and Supplemental Table 1-A)** at all DRG levels. Having determined that paclitaxel affects nociceptors in all DRGs, we pooled samples from 4 mice (2 male and 2 female) per treatment, per biological replicate for high-throughput sequencing (TRAP-seq). Subsequent experiments were also carried out in males and females **(Supplemental Table 2)** and no differences were observed between sexes. On day 10 after the start of paclitaxel or vehicle treatment, we collected eGFP-L10a IPs from Nav1.8-TRAP mice for RNA sequencing. To gain insight into the specificity of the Nav1.8-TRAP approach, we analyzed a subset of genes that are known to be enriched in different cell populations in the DRG. This analysis demonstrated that TRAP-seq allowed the isolation of translating mRNAs in a cell-type specific manner. Indeed, neuronal mRNAs such as *Calca* (CGRPα), *Calcb* (CGRPβ) or *Scn10a* (Nav1.8) were enriched in TRAP fractions whereas glial markers (*Aldh1l1*, *Gfap)* and endothelial specific genes (*Cldn5, Esam)* were depleted. **(Fig. 1D)**.

From all biological replicates we sequenced input mRNA, equivalent to the DRG transcriptome, and TRAP mRNAs associated with translating ribosomes in the Nav1.8 subset of DRG neurons. This approach allowed us to make comparisons of transcriptional and translational changes in response to neuropathic pain. Gene expression values (TPM) were normalized to the 90^th^ percentile for each biological replicate and the empirical probability density function (PDF) of the normalized expression level (upper decile (ud)TPM) was plotted for the input and TRAP fractions **(Fig. S3A,B).** The PDF function displayed 2 peaks **(Fig. S3A,B)** and the inflexion point was used to set the threshold expression values according to the sequencing depth. After further filtering, based on consistent expression among biological replicates, we included a total of 8270 genes in the final analysis.

We then plotted the cumulative frequency distribution as a function of the log 2-fold change for each of these genes in vehicle-versus paclitaxel-treated biological replicates and the 95^th^ percentile was used to set the threshold fold change values for the input and TRAP fractions **(Fig. S3D)**. Input transcriptome analysis revealed that only 4 genes were up-regulated (*Dpep2, Car3, Iah1* and *Arhgef4*) and 1 was down-regulated (*Plcb3*) in the DRG by paclitaxel treatment (**Fig. 1E** and **Supplemental Table 3**) suggesting that at this time point there are relatively few transcription changes induced by paclitaxel treatment in the whole DRG. In stark contrast, and consistent with the notion that transcriptional and translational changes are frequently decoupled (*9*), we detected 230 genes up-regulated and 222 down regulated by paclitaxel treatment in the TRAP-seq data set **(Fig. 1E and F** and **Supplemental Table 4 and 5)**. The difference in the number of genes differentially expressed in the input versus the TRAP-seq is unlikely due to variability between samples because strong correlation coefficients were observed between biological replicates **(Fig. S4A)** and principal component analysis did not distinguish between treatment conditions in input samples whereas robust differences were found in the TRAP-seq samples **(Fig. S4B,C and D)**.

Combining this dataset with single cell RNA sequencing from existing data sources (*10, 11*) allowed us to infer translation efficiencies (TEs) for all mRNAs translated in Nav1.8 neurons creating a rich resource for understanding steady state translation of mRNAs expressed in nociceptors (https://www.utdallas.edu/bbs/painneurosciencelab/sensoryomics/cipntrap/browse.html). We examined families of genes for any systematic differences in TEs for mRNAs expressed in Nav1.8-positive nociceptors (e.g. ion channels, G-protein coupled receptors and kinases). Interestingly, we observed that ion channels and GPCR tend to show higher TEs compared to other gene families such as kinases, RNA-binding proteins or transcription factors **(Fig. S5A).** With paclitaxel treatment we found a clear increase in TE for *Trpv1*, which has been shown to be up-regulated after paclitaxel (*12*), and *P2rx2,* and a decrease in TE for *Chrna6*. These latter two genes interact such that *Chrna6* inhibits the pronociceptive activity of *P2rx2* (*13*) suggesting a reciprocal translational regulation event that would be expect to enhance pain sensitivity **(Fig. S5B)**.

Having established alterations in the nociceptor translatome with paclitaxel treatment, we then sought to examine whether there were motifs in the 5’ untranslated regions (UTRs) of mRNAs up-regulated after CIPN. We found an enrichment for 3 motifs, among them a terminal oligopyrimidine tract (TOP)-like element which is associated with mTORC1-mediated translation control (*14*) **(Fig. S6)**. Another enriched motif was a G-quartet-like sequence suggesting translation control by the RNA helicase eIF4A (*15*) **(Fig. S6)**. Interestingly, *eIF4a3* was highly translated after CIPN (Supplemental Table 4). The final motif was an A/G rich sequence with no known function.

We then investigated what classes of genes were significantly enriched in the up and down regulated mRNA TRAP-seq dataset using the EnrichR tool (*16*). First, validating that this approach can identify pathological changes in nociceptors that have already been linked to CIPN (*17*), reactive oxygen species and mitochondrial dysregulation signatures were significantly enriched in the paclitaxel-treated TRAP-seq dataset. Likewise, several translationally up-regulated mRNAs have already been identified in the CIPN literature using independent methods, such as caspase 6 (*18*). We also found a significant enrichment of genes involved in the regulation of cap-dependent translation, the lysosomal complex and mTORC1 signaling **(Fig. 1G)**. Closer examination of the up-regulated and down-regulated mRNA lists for TRAP-seq with EnrichR analysis led us to further investigate a network of mTORC1 signaling-associated proteins that we hypothesized are critical for pathology causing CIPN.

We confirmed changes in steady state protein levels for a panel of targets identified in the TRAP-seq dataset with increased translation efficiency with paclitaxel treatment **(Fig. 2A-B)** including mTOR **(Fig. 2B)**, and Esyt1 **(Fig. S7A-C).** The concordance between measurement indicates that TRAP-seq can reliably predict increases in protein expression in DRG after CIPN. Based on increased mTOR protein levels we also examined downstream and associated targets of mTOR where we noted increases in p-4EBP1 expression **(Fig. 2B)** but no differences in phosphorylation or total protein levels for ERK, AKT or, surprisingly, ribosomal protein S6 **(Fig. S8A-B)**. We also found that eIF4E phosphorylation was increased with paclitaxel treatment **(Fig. 2B)**. These findings suggest a selective increase in signaling via a subset of proteins that control translation of genes involved in neuronal plasticity (*19, 20*). To test this hypothesis, we asked if mTOR inhibition or genetic loss of the phosphorylation site for mitogen activated protein kinase interacting kinase (MNK) on the cap-binding protein eIF4E (eIF4E^S209A^) would impair CIPN.

**Figure 2.**
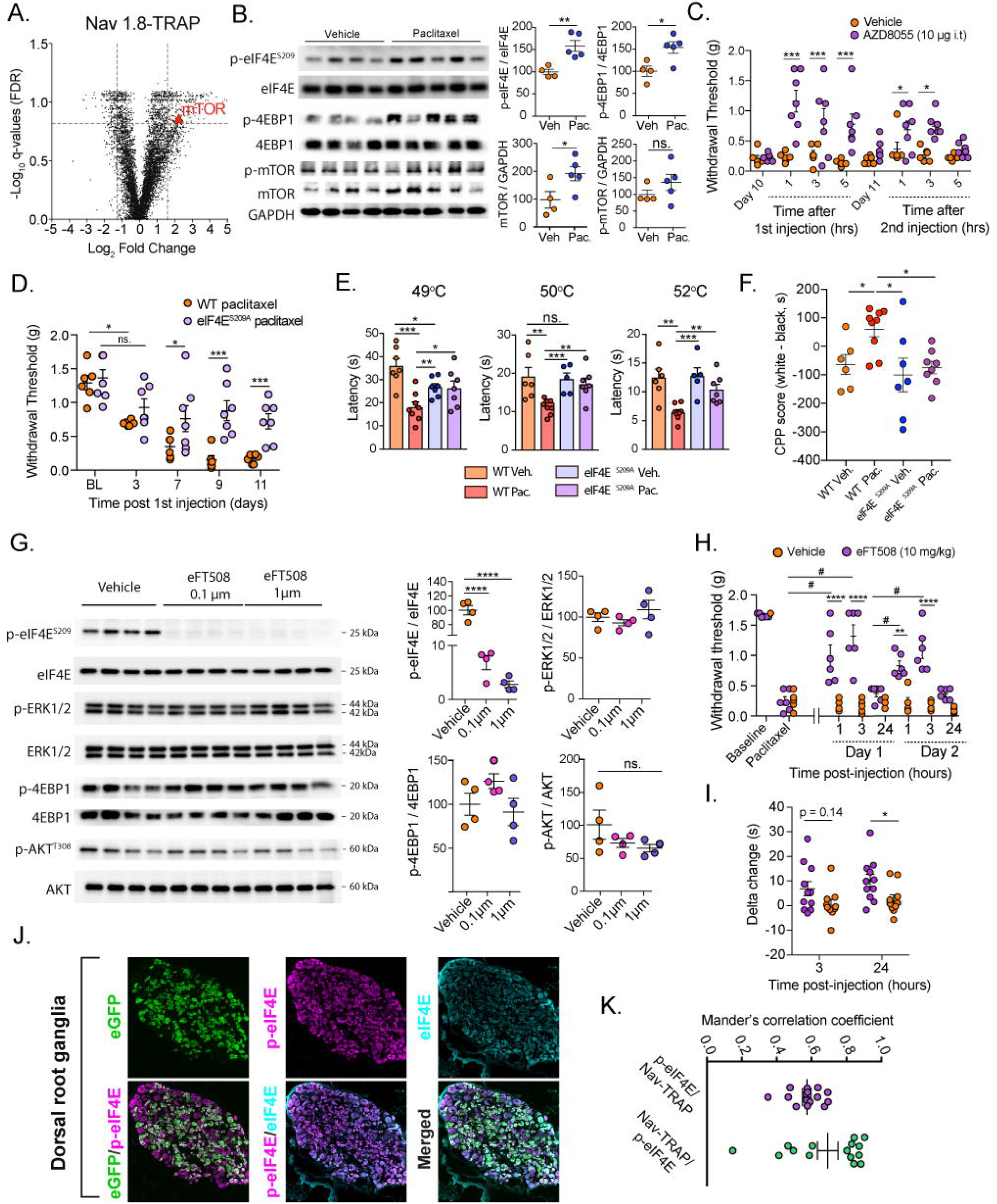
Enhanced translation regulation signaling in nociceptors is responsible for evoked and spontaneous pain in paclitaxel-induced neuropathic pain. **(A)** Volcano plot showing an increase in *mTOR* mRNA translation in the DRGs after CIPN.**(B)** Immunoblotting shows an up-regulation of p-eIF4E, p-4EBP1,and mTOR after paclitaxel treatment (p-eiF4E: Veh = 100 ± 5.39, Pac = 158 ± 10.57, **p = 0.0026, n = 4-5; p-4EBP1: Veh = 100 ± 9.8, Pac = 152 ± 11.27, *p = 0.012, n = 4-5; p-mTOR: Veh = 100 ± 10.8, Pac = 131 ± 20.27, p = 0.23, n = 4-5; mTOR: Veh = 100 ± 24.54, Pac = 195 ± 25.27, *p = 0.025, n = 4-5). **(C)** Intrathecal injection of the mTOR kinase inhibitor AZD8055 relieves= mechanical allodynia compared to vehicle-treated animals after the first injection (Two way ANOVA: F_(7,88)_ = 3.93; post-hoc Bonferoni: at 1hr: ****p<0.0001, 3hrs: *p = 0.043, 5hrs: **p = 0.0059) but no significant effects were observed after the second injection. **(D)** eIF4E^S209A^ mice show a deficit in development of paclitaxel-induced mechanical allodynia compared to WT animals. Two way ANOVA: F_(4,44)_ = 3.14, p = 0.023 with a post-hoc test: at day 7: *p = 0.045; at day 9: ***p = 0.0003; at day 11: **p = 0.0081 eIF4E^S209A^ vs WT, n = 7. **(E)** eIF4E^S209A^ mice also exhibit a deficit in paclitaxel-induced thermal hyperalgesia compared to WT animals at 49°C (One-way ANOVA: F = 8.39, p = 0.005; p°st-h°c B°nfer°ni; Veh WT vs Pac WT:** p = 0.005, Veh WT vs Pac eIF4E^S209A^: *p = 0.071, Veh WT vs Veh eIF4E^S209A^: *p = 0.054), at 50°C (One-way ANOVA: F =6.23, p = 0.0032; post-hoc Bonferoni; Veh WT vs Pac WT: **p = 0.0044, Veh WT vs Pac eIF4E^S209A^: p > 0.99, Veh WT vs Veh eIF4E^S209A^: p > 0.99, Pac WT vs Pac eIF4E^S209A^: *p = 0.024) and 52°C (One-way ANOVA: F =7.78, p = 0.0010; post-hoc Bonferoni; Veh WT vs Pac WT: **p = 0.0021, Veh WT vs Pac eIF4E^S209A^: p > 0.99, Veh WT vs Veh eIF4E^S209A^: p > 0.99, Pac WT vs Pac eIF4E^S209A^: *p = 0.045). **(F)** WT paclitaxel-treated mice show signs of ongoing pain in the CPP paradigm compared to WT vehicle (One way ANOVA: F= 4.36, p = 0.012, post-hoc Bonferoni Veh WT vs Pac WT: *p = 0.048) and eIF4E^S209A^ mice treated with paclitaxel, which did not show signs of spontaneous pain (One way ANOVA: F= 4.36, p = 0.012, post-hoc Bonferoni Veh WT vs Pac WT: *p = 0.019). **(G)** eFT508 significantly reduced the level of eIF4E phosphorylation at 0.1 and 1 μM after 1 hr (One way ANOVA: F = 205.2, p<0.0001; post-hoc Tukey: **** p< 0.0001), without affecting other signaling pathways in cultured mouse DRG neurons. **(H)** Acute treatment with eFT08 relieves mechanical allodynia compared to vehicle after the 1^st^ (Two way ANOVA: F(7,63) = 14.53, post-hoc Bonferoni at 1 and 3 hrs: ****p<0.0001) and the 2nd injection (Two way ANOVA: F_(7,63)_ = 14.53, post-hoc Bonferoni at 1hr: *** p<0.001 and 3hrs: ****p<0.0001). **(I)** Acute treatment with eFT508 did not significantly attenuate thermal hyperalgesia at 3 hrs (Two way ANOVA: F_(1,38)_ = 8.46, post-hoc Sidak: p = 0.14) but a significant effect was observed at 24 hrs (Two way ANOVA: F_(1,38)_ = 8.46, post-hoc Sidak: *p < 0.05). **J)** and **K)** p-eIF4E is highly expressed in Nav1.8-positive DRG neurons and approximatively 80% of the eGFPe signal co-localizes with p-eIF4E.

We used the ATP competitive mTORC1/2 inhibitor AZD8055 to examine the influence of mTOR inhibition on paclitaxel-induced pain. While a robust reduction in mechanical hypersensitivity occurred with the first intrathecal injection of AZD8055, the magnitude of this effect was decreased upon subsequent treatments **(Fig. 2C)**. We attribute this rapid onset, tolerance-like effect to the known phenomena of feedback signaling in the mTOR pathway (*21, 22*). We then turned to a genetic approach to examine disrupted MNK-eIF4E signaling. eIF4E^S209A^ mice treated with paclitaxel showed decreased mechanical and thermal hypersensitivity compared to their WT littermates **(Fig. 2D-E).** While this suggests decreased CIPN pain in the absence of eIF4E phosphorylation, we determined this empirically using the conditioned place preference (CPP) paradigm (*23*) with retigabine as the conditioning agent, as has been described previously (*24*). Importantly, WT mice had a clear preference for the retigabine-paired chamber demonstrating relief of spontaneous pain. On the other hand, *eIF4E*^*S209A*^ mice showed no preference for the retigabine-paired chamber. This indicates an absence of spontaneous pain in these mice **(Fig. 2F)**.

Given the strong behavioral phenotype of eIF4E^S209A^ mice in CIPN, we turned to a clinical candidate drug, eFT508 (*25*). eFT508 is a potent MNK1/2 inhibitor and blocks eIF4E phosphorylation *in vitro* and *in vivo.* We used this compound to examine whether targeted blockade of the MNK-eIF4E signalling axis inhibits and/or reverses paclitaxel-induced pain. We first tested the potency of eFT508 in cultured DRGs neurons. Treatment with eFT508 reduced eIF4E phosphorylation by 90% without affecting ERK, 4EBP or AKT phosphorylation or expression **(Fig. 2G and Fig. S9).** eFT508 did not reduce bulk translation **(Fig. S10A-B).** This parallels previous findings that Mnk1/2 controls the translation of a specific subset of mRNAs (*19, 26*). Paclitaxel-treated mice given eFT508 after full development of CIPN showed a dose-related reversal of mechanical and thermal hypersensitivity **(Fig. 2H-I)**. Also, the animals did not show sign of tolerance with multiple treatments and the effect of eFT508 was completely absent in eIF4E^S209A^ mice demonstrating the specificity of te drug **(Fig. S11).** Consistent with an action on nociceptors, 80% of eGFP-L10a+ neurons exhibited strong p-eIF4E signal **(Fig. 2J-K).** No signal for the p-eIF4E antibody was observed in the eIF4E^S209A^ mice, indicative of the specificity of the antibody **(Fig. S12)**.

Our findings suggest a link between increased translation of mTOR complex proteins and MNK-mediated eIF4E phosphorylation as a causative factor in paclitaxel-induced neuropathy. However, mechanisms through which eIF4E phosphorylation might control mTOR signaling are not established. A possible connection between these parallel pathways (*27*) is reciprocal translational regulation of mRNAs that encode signaling factors of the RagA-Ragulator complex mediated by MNK-dependent eIF4E phosphorylation. The RagA-Ragulator complex controls mTORC1 activity in response to amino acid flux across the lysosomal membrane (*28, 29*) and requires activity of the RagA, but not the RagB, GTPase (*30, 31*). RagA exists in a complex with RagC while RagB forms a complex with RagD (*30, 31*). Our TRAP-seq comparison demonstrates that genes corresponding to the components of the RagA-Ragulator-mTORC1 complex are preferentially translated after paclitaxel treatment **(Fig. 3A).** In particular, there was a 3.5 fold increase in the number of normalized reads mapping to the *Rraga* mRNA, encoding the RagA GTPase amino acid sensor for mTORC1 **(Fig. 3B-C)**.

**Figure 3.**
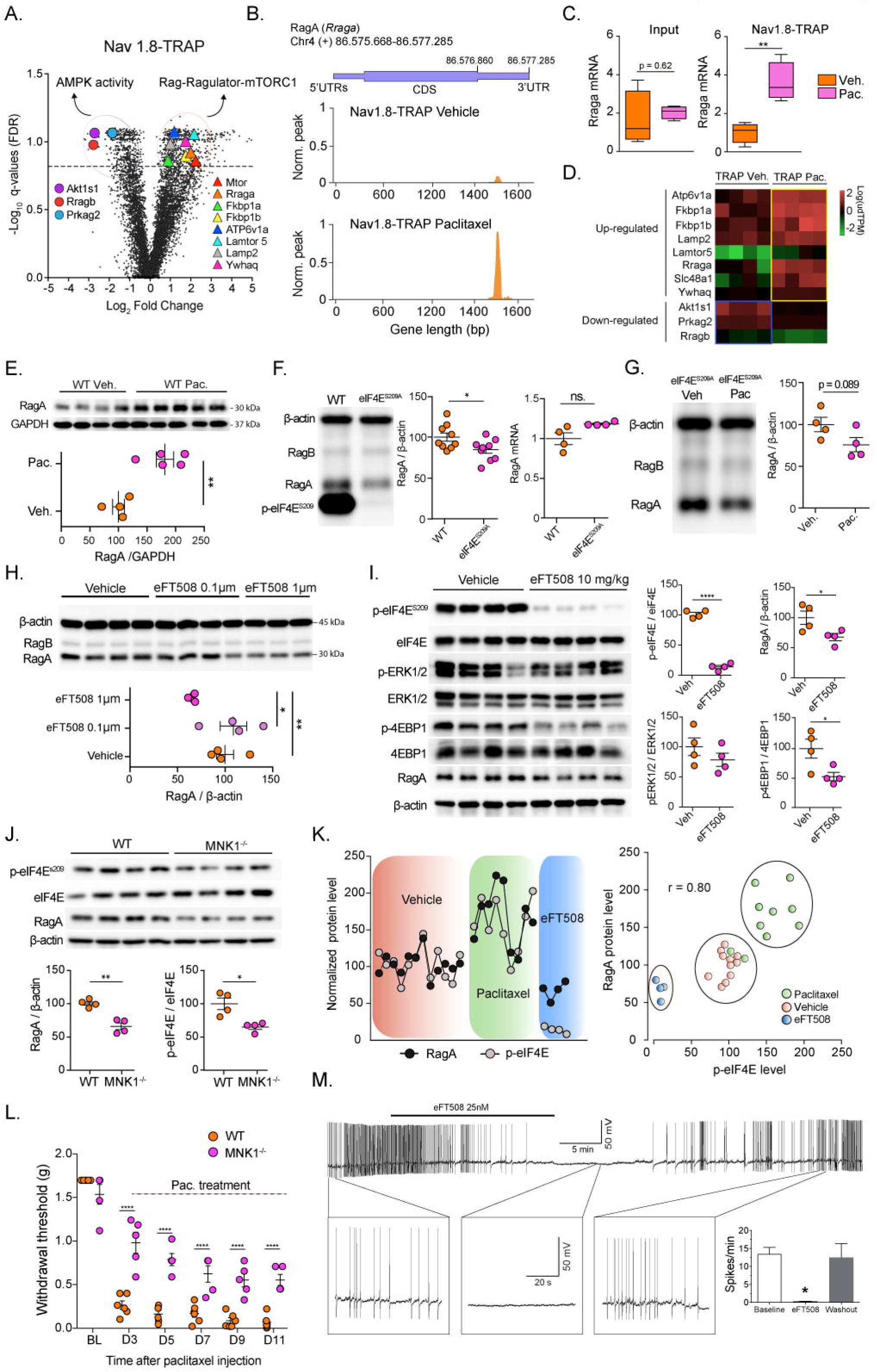
Mnk1-eIF4E signaling controls the translation of RagA GTPase driving CIPN. **(A)** Volcano plot shows an increase in translation efficiency for mRNAs encoding proteins associated with the Rag-Ragulator-mTORC1 complex. **(B)** Distribution of the normalized reads on the RagA gene (Rraga) showing a prominent peak in the TRAP fraction after CIPN compared to Veh while no changes are observed in the Input fraction. **(C)** Normalized TPM for the Rraga mRNA shows no differences in the input (RagA: Veh = 1.65 ± 0.71, Pac: 2.04 ± 0.17 p = 0.62) while a significant increase is observes in the TRAP fraction (RagA: Veh = 1.004 ± 0.26, Pac: 3.6 ± 0.51 p = 0.0043). **(D)** Heatmap showing increased or decreased translation of the mRNAs encoding proteins of the Ragulator-RagA-mTORC1 complex after CIPN. **(E)** Immunoblot showing an increase in RagA protein levels in DRGs after CIPN compared to Veh (RagA: Veh = 100 ± 10.33, Pac = 181 ± 15.35, **p = 0.0043, n = 4-5). **(F)** Immunoblot showing a decrease in RagA protein levels in the eIF4E^S209A^ mice compared to WT (RagA: Veh = 100 ± 5.05, Pac = 84.6 ± 4.32, *p = 0.035, n = 9) while no differences were observed in mRNA levels (RagA: Veh = 1.007 ± 0.067, Pac = 1.176 ± 0.01, p = 0.084, n = 4) **(G)** Immunoblot showing no differences in the DRG level of RagA protein between eIF4E^S209A^ vehicle-and paclitaxel-treated mice (RagA: Veh = 100 ± 8.567, Pac = 75 ± 8.46, *p = 0.095, n = 4). **(H)** Immunoblot showing a decrease in RagA protein level after treatment with eFT508 at 1 μM for 1 hr (One way ANOVA: F= 4.55, p = 0.040, post-hoc Sidak Veh WT vs: 1 μM *p = 0.042). **(I)** Immunoblot showing a significant decrease in the level of p-eIF4E and RagA after eFT508 treatment in DRGs *in vivo* (p-eIF4E: Veh = 100 ± 2.33, eFT508: 10 ± 2.43, ****p<0.0001, n = 4; p4EBP1: Veh = 100 ± 16.03, eFT508 = 53.12 ± 7.03, *p<0.05; RagA: Veh = 100 ± 11.03, eFT508: 67 ± 6.02, *p<0. 042, n = 4 **(J)** Immunoblot showing a significant decrease in the level of p-eIF4E (Veh = 100 ± 8.63, MNK1^-/-^: 64.98 ± 3.63, **p<0.0094, n = 4) and RagA (Veh = 100 ± 2.54, MNK1^-/-^: 67 ± 5.70, *p<0. 0010, n = 4) in *MNK1*^-/-^mice compared to WT. **(K)** Co-expression level of RagA and p-eIF4E in vehicle-, paclitaxel-and eFT508-treated mice shows high correlation between DRG samples. **(L)** *MNK1*^-/-^ show a deficit in development of paclitaxel-induced mechanical allodynia compared to WT animals ((Two way ANOVA: F_(5,45)_ = 13.45, p < 0.0001 with a post-hoc test: at day 3, 5, 7, 9 ****p<0.0001 *MNK1*^-/-^ vs WT, n = 7). **(M)** DRG neurons taken from mice treated with paclitaxel show spontaneous activity that is suppressed by eFT508 in a reversible fashion.

Other components of the RagA-Ragulator-mTORC1 complex including *Lamtor5, Lamp2, Atp6v1a, Fkbp1a, Fkbp1b, Ywhaq* and *Mtor* also showed a consistent increase in translation with paclitaxel treatment **(Fig. 3B-C)**. Interestingly, known negative regulators of mTORC1 including *Akt1s1*, encoding PRAS40, and the γ2 subunit of AMPK (*Prkag2*) showed decrease translation efficiency **(Fig. 3A and D)**. These findings indicate a fundamental change in the mTORC1 signaling network in Nav1.8 expressing neurons in CIPN that could be indicative of altered protein biosynthesis or related to other functions of mTORC1, such as lipid homeostasis (*32*). A component of this latter pathway, *Soat2*, was also selectively translated with paclitaxel and can be inhibited by avasimibe (*33*). Repeated treatment with this drug had no effect on paclitaxel-induced pain **(Fig. S13)**. Given the behavioral observations with eIF4E^S209A^ mice, the effect of eFT508 in CIPN and negative findings with avasimibe, we further focused on translation regulation mechanisms as the key causative factor in paclitaxel-induced neuropathic pain.

We validated that RagA protein levels were increased in DRG by paclitaxel treatment **(Fig. 3E)**. We also found that RagA protein expression was significantly lower in *eIF4E*^*S209A*^ mice compared to WT mice, although mRNA levels were equal in the two strains of mice **(Fig. 3F)**. These findings suggested that eIF4E phosphorylation controls RagA translation in CIPN. To test this, we treated eIF4E^S209A^ mice with paclitaxel and failed to observe an effect from paclitaxel on RagA protein in DRGs from these mice **(Fig. 3G)**. Moreover, the MNK1/2 inhibitor eFT508 decreased RagA protein levels both in cultured DRG neurons **(Fig. 3H)** and *in vivo* **(Fig. 3I)**. 4EBP1 phosphorylation was also decreased in the DRG of eFT508-treated mice linking altered RagA translation to downstream mTORC1 activity **(Fig. 3I)**. Interestingly, MNK1/2 inhibition also decreased the protein expression level of Lamtor5 in DRG neurons **(Fig. S14).**

Two MNK isoforms are known to phosphorylate eIF4E, however, MNK2 shows constitutive kinase activity while MNK1 is induced by cellular stimuli (*34*). We reasoned that MNK1-mediated phosphorylation of eIF4E might control RagA protein expression and regulate paclitaxel-induced CIPN. *MNK1*^-/-^ mice showed a 40% reduction in RagA protein levels in DRG compared to WT and a significant reduction in eIF4E phosphorylation while upstream signaling pathways were unchanged **(Fig. 3J, Fig. S15A).** When we co-examined eIF4E phosphorylation with RagA expression in mouse DRG we observed correlated expression for RagA and p-eIF4E. This was increased by paclitaxel treatment and strongly suppressed by eFT508 with an 80% correlation coefficient indicating that increases or decreases in eiF4E phosphorylation can reliably influence the level of RagA translation **(Fig. 3K)**. We then assessed paclitaxel-induced CIPN in *MNK1*^-/-^ mice and observed a profound deficit in the development of neuropathic pain **(Fig. 3L)** consistent with observations in *eIF4E*^*S209A*^ mice. Finally, we examined whether transient MNK1/2 inhibition could decrease spontaneous activity in DRG neurons taken from paclitaxel treated mice. These neurons showed robust spontaneous activity that was almost completely suppressed by eFT508 treatment within 20 min **(Fig 3M and Supplemental Table 1-B)**. This effect was reversible indicating that eFT508 is not toxic to DRG neurons *in vitro*.

We used nociceptor translational profiling in a mouse model of neuropathic pain to provide a rich resource for probing translational control in pain plasticity and reveal high quality targets for the potential generation of disease modifying therapies for a currently intractable disease. Our work demonstrates dysregulation of the mTORC1 signaling network in CIPN that reveals a complex interplay between eIF4E-mediated translation control of RagA and mTORC1 function **(Fig. S15B)**. Our experiments support a model wherein MNK1-mediated phosphorylation of eIF4E controls the translation of RagA mRNA creating a link between these distinct signaling pathways. In the context of CIPN, increased eIF4E phosphorylation leads to enhanced RagA protein levels driving mTORC1 activation and neuropathic pain. Our findings demonstrate that MNK and eIF4E phosphorylation can be targeted genetically or with eFT508, a drug in late phase clinical trials, to prevent or reverse CIPN pain. These discoveries can be used to enhance the efficacy of cancer chemotherapy treatment by reducing the primary dose-limiting side effect of chemotherapy and can improve the quality of life of people who suffer from CIPN.

## Acknowledgements

This work was supported by NIH grants R01NS065926 (TJP), R01NS098826 (TJP and GD), R01CA200263 (PMD) and R01NS100788 (ZTC), The University of Texas STARS program (TJP and GD), The H.E.B Professorship in Cancer Research (PMD) and a postdoctoral CONACYT fellowship program (PBI).

## Author contributions

S.M., Z.T.C. and T.J.P. conceived of the project. S.M., P.R.R., P.M.D., A.K., N.S., K.R.W., Z.T.C., G.D. and T.J.P. designed experiments. S.M, J.K.M. and L.T.F. performed protein expression and biochemical experiments, with assistance from P.B.I. S.M., P.R.R., A.W. and Z.T.C. analyzed sequencing data. S.M., J.K.M. and A.A. did behavioral experiments. Y.L. and R.N. performed electrophysiological recordings. S.M., Z.T.C. and T.J.P. wrote the manuscript. All authors approved the manuscript.

## Competing interests

The authors declare no competing interests.

## Data and materials availability

Raw RNA sequencing data are available through GEO. Transgenic mice are available through Jackson Laboratories. All raw data and code is available upon request.

## Supplementary materials

Materials and Methods

Figs. S1 to S15

Table S1 to S7

References (1–10)

